# Epigenetic contributions to hemisphere asymmetry in healthy brain, aging, and Parkinson’s disease

**DOI:** 10.1101/794255

**Authors:** Peipei Li, Elizabeth Ensink, Sean Lang, Lee Marshall, Meghan Schilthuis, Jared Lamp, Irving Vega, Viviane Labrie

## Abstract

Hemispheric asymmetry in neuronal processes is a fundamental feature of the human brain and drives symptom lateralization in Parkinson’s disease (PD), but its molecular determinants are unknown. Here, we determine epigenetic differences involved in hemispheric asymmetry in the healthy and the PD brain. Neurons of the healthy brain exhibit numerous hemispheric differences in DNA methylation, which affect genes implicated in neurodegenerative diseases. In PD patients, hemispheric asymmetry in DNA methylation is even greater and involves many PD risk genes. The lateralization of clinical PD symptoms involves epigenetic, transcriptional, and proteomic differences across hemispheres that affect neurodevelopment, immune activation, and synaptic transmission. During aging, healthy neurons show a progressive loss of hemispheric asymmetry in the epigenome, which is amplified in PD. For PD patients, a long disease course is associated with greater hemispheric asymmetry in neuronal epigenomes than a short disease course. Hemispheric differences in epigenetic gene regulation are prevalent in neurons and may affect the progression and symptoms of PD.

## Introduction

Parkinson’s disease (PD) is a severe, irreversible neurodegenerative disease involving motor symptoms that are unilateral at onset for over 85% of patients^1–3^. The lateralization of motor symptoms results from an asymmetric pattern of neurodegeneration in the brain^4–8^. PD patients have hemispheric asymmetry in neuronal dysfunction in both the nigrostriatal system and cortical brain structures^4–8^, such that the hemisphere contralateral to PD motor symptom predominance shows greater neuronal and synaptic dysfunction than the ipsilateral side^4–8^. Right/left brain asymmetries in PD appear early, in preclinical stages^4, 8, 9^. Though PD symptoms eventually affect both body sides as the disease progresses, clinical asymmetry remains directionally stable and detectable even at advanced disease stages^10, 11^. Furthermore, asymmetric motor presentation is linked to the rate of disease progression^12–14^. Cognitive symptoms also differ between subgroups of lateralized PD patients (i.e., visuospatial tasks, language, verbal memory, and psychosis differ between patients with left *vs.* right motor symptom predominance)^14–19^. Despite the prevalence of asymmetric brain changes in PD and its relevance to disease progression and clinical manifestations, the factors rendering neurons more vulnerable to degeneration in one hemisphere over the other are unknown.

Epigenetic regulation represents a mechanism through which genetic, environmental, and aging risk factors could plausibly trigger hemispheric differences in neuronal and synaptic loss. Epigenetic marks like DNA methylation enable dynamic regulation of gene expression throughout the life of a neuron^20, 21^. In the brain, divergent DNA methylation signatures facilitate the functional specialization of neurons and brain subregions^22–24^. During early development, hemispheric asymmetry in DNA methylation contributes to the lateralization of nervous system organization, which affects hemisphere dominance for handedness, cognitive processes, and language^25–27^. Furthermore, DNA methylation status governs the activity of gene regulatory elements such as enhancers and promoters, which affect the establishment of left–right asymmetries in various tissues^28–30^. Hence, epigenetic variation may influence asymmetrical gene expression patterns in the brain, which if pathogenic could contribute to PD.

In post-mitotic neurons, disruption of DNA methylation induces lasting changes in synaptic architecture and cellular signaling that can promote neurodegenerative processes^31–33^. Genome-wide studies have identified abnormalities in DNA methylation in the PD brain^34–37^. In addition, several studies of the PD brain have demonstrated a loss of DNA methylation at the α-synuclein gene promoter, which may contribute to elevated α-synuclein expression^38–40^; a major PD risk factor and the main component of Lewy pathology in this disease. Furthermore, aging is the strongest risk factor for PD, and epigenetic changes contribute to aging processes^20, 41, 42^. Genes linked to neurodegeneration exhibit epigenetic changes with age^43^, and accelerated epigenetic aging is observed in PD^44^. A hemispheric imbalance in the accumulation of DNA methylation abnormalities affecting genes involved in disease pathophysiology may explain the lateralization of clinical symptoms in PD, though this has yet to be examined.

Here, we identify DNA methylation patterns involved in hemispheric asymmetry in the healthy (neurotypical) and PD brain. DNA methylation was fine-mapped at gene regulatory elements, enhancers and promoters, genome-wide, in isolated neurons of the prefrontal cortex of PD patients and controls. In two independent cohorts, we find that neurons of PD patients have extensive hemispheric asymmetry in DNA methylation, exceeding that of control individuals. In particular, regulatory elements of PD risk genes (identified in genetic studies) show prominent epigenetic asymmetry in PD patients. Inter-hemispheric differences in the epigenome of PD patients closely associates with symptom lateralization, such that epigenetic changes are most apparent in the hemisphere matched to the predominant side of symptom presentation. Furthermore, hemispheric asymmetry in DNA methylation patterns is linked to genes with transcriptomic and proteomic differences between hemispheres. In aging, there is a progressive loss of hemispheric asymmetry in the epigenomes of the healthy brain that is also evident in PD. For PD patients, epigenetic asymmetry between hemispheres is associated with differences in disease progression, as PD patients with a long disease course have more hemispheric asymmetry. Together, our results support that hemispheric asymmetry in PD involves the disrupted epigenetic regulation of genes, which may enable asymmetric neuronal dysfunction and the contralateral predominance of PD symptoms.

## Results

### Hemispheric asymmetry involves aberrant DNA methylation in PD neurons

To determine whether there are hemispheric differences in DNA methylation that could impact neuronal function in the healthy (neurotypical) and PD brain, we comprehensively fine-mapped DNA methylation in neurons isolated from either the left or right prefrontal cortex of PD patients and controls (n = 57 and 48 individuals, respectively; Supplementary Data 1). Neuronal nuclei from hemispheres were isolated by an established antibody- and flow cytometry-based approach^32, 45^ (Supplementary Fig 1). DNA methylation was profiled at all brain enhancers and promoters across the genome, including active and poised/bivalent elements, as defined by the NIH Roadmap Epigenomics Project (ChromHMM 18-state model). Genome-wide mapping of DNA methylation at enhancers and promoters was performed with a targeted bisulfite sequencing strategy, known as the bisulfite padlock probe approach. The padlock probe library consisted of 59,009 probes targeting 35,288 regulatory elements (Supplementary Data 2). In PD and control neurons, we investigated a total of 623,041 modified cytosines, of which 105,238 were CpG and 517,803 were CpH sites (Supplementary Fig. 2).

**Fig. 1.**
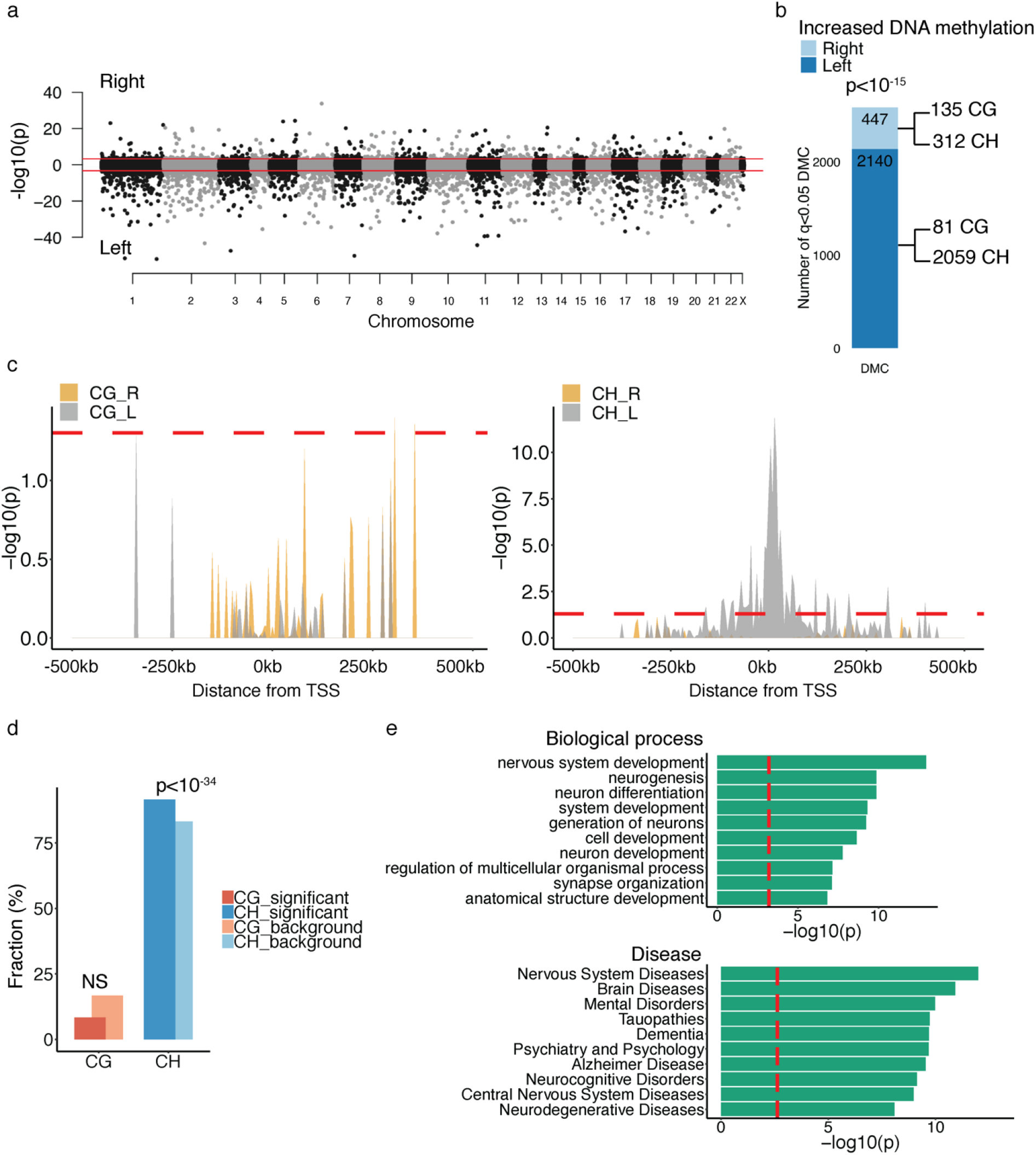
Hemispheric asymmetry in DNA methylation in neurons of the healthy human brain. (**a**) Manhattan plot showing DNA methylation differences between neurons of the left and right hemisphere of the healthy brain, after controlling for age, sex, postmortem interval, and neuronal subtype proportion. DNA methylation was mapped at enhancers and promoters of neurons from the healthy prefrontal cortex (n = 25 left hemispheres, 23 right). −log10(p) refers to the significance of differentially methylated cytosines, with the sign corresponding to the hemisphere side exhibiting higher DNA methylation levels. Threshold for genome-wide significance (red line) is *q* < 0.05, as determined by robust linear regression with contrasts. (**b**) Bar plot showing the distribution of differentially methylated cytosines (DMC) across hemispheres. The number of significant cytosine sites that show greater DNA methylation levels on the left or right hemisphere is shown. The *p*-value represents the enrichment of cytosines with increased DNA methylation in the left hemisphere, by Fisher’s exact test. CpG and CpH site contribution to differential methylation in each hemisphere is shown. (**c**) Genomic location of DNA methylation changes involved in hemispheric asymmetry in neurons of the healthy brain. The location of CpG (left panel) and CpH (right panel) sites with increased DNA methylation in the left or right hemisphere is shown. Location enrichment was determined by hypergeometric test of differentially methylated cytosines compared to background cytosines (5 kb bins), and relative to the distance from the nearest transcription start site. Red dashed line, *p* < 0.05. (**d**) CpG and CpH involvement in inter-hemispheric DNA methylation differences. The percent number of significantly altered CpG or CpH sites differing between hemispheres (relative to background) is shown. The enrichment of cytosine context involved in hemispheric asymmetry was determined by hypergeometric test. (**e**) Top biological processes and disease pathways of genes affected by hemispheric asymmetry in DNA methylation. Pathway analysis was done by MetaCore. Threshold for significance (red dashed line) is *q* < 0.05.

We examined whether there were left–right hemispheric differences in the epigenome of healthy neurons, and whether hemispheric differences were greater in PD neurons than in control neurons. Our analysis examined individual cytosine sites (CpGs and CpHs) for hemispheric asymmetry in control and PD neurons, and adjusted for age, sex, post-mortem interval, and neuronal subtype proportion. Neuronal subtype proportion refers to the proportion of glutamatergic to GABAergic neurons, as determined by neuronal subtype deconvolution. The PD prefrontal cortex did not exhibit a hemisphere-specific loss in subtypes of glutamatergic and GABAergic neurons (Supplementary Data 3).

Neurons of the healthy human brain exhibited a prominent hemispheric asymmetry in DNA methylation (n = 25 left and 23 right hemisphere; Fig. 1a). Inter-hemispheric differences in DNA methylation occurred at 2,587 cytosine sites at enhancers and promoters in healthy neurons (*q* < 0.05, robust linear regression with contrasts; Fig. 1a; Supplementary Data 4). In particular, the left hemisphere had higher DNA methylation levels than the right hemisphere (82.7% of significant cytosines had more DNA methylation in the left hemisphere; *p* < 10^-15^, Fisher’s exact test; Fig. 1b), which was largely due to differential CpH methylation (Fig. 1b and 1c). Indeed, CpH methylation, a silencing epigenetic mark at enhancers and promoters^21^, was the primary epigenetic contributor to hemispheric asymmetry (2,371 CpHs and 216 CpGs, *p* < 10^-34^ for CpH enrichment, hypergeometric test; Fig. 1d). In healthy neurons, the hemispheric asymmetry of DNA methylation most often affected cis-acting regulatory elements, on average within 47.2 ± 1.4 kb of transcription start sites (average DNA methylation change 6.09 ± 0.39% at CpGs and 1.34 ± 0.05% at CpHs; Fig. 1c).

We then determined the gene targets of the enhancers and promoters that showed hemispheric asymmetry in their epigenome. We analyzed 3D chromatin interactions (Hi-C data) in the human prefrontal cortex^46^, and found 1,995 genes targeted by enhancers and promoters with hemispheric asymmetry. To further capture proximal interactions, we used an *in silico* cis-regulatory element prediction tool^47^. In total, we found 3,793 genes having hemispheric differences in the epigenetic regulation of their cis-regulatory elements (Supplementary Data 5). Pathway analysis revealed that genes with hemispheric asymmetry in epigenetic regulation were primarily involved in neurodevelopment, synaptic organization, and neurodegenerative illnesses (*q* < 0.05, hypergeometric test; Fig. 1e). These findings demonstrate that hemispheric asymmetry in neuronal epigenomes is prevalent in the healthy brain and may lead to inter-hemispheric differences in vulnerability to neurodegeneration.

We next determined that PD patient neurons have substantially more hemispheric asymmetry in DNA methylation than control neurons (n = 23 PD-left hemisphere, 34 PD-right, 25 control-left, 23 control-right). There were 6,207 cytosine sites exhibiting left *vs.* right hemispheric asymmetry in DNA methylation at enhancers and promoters (*q* < 0.05, robust linear regression with contrasts; Fig. 2a; Supplementary Data 4), of which 3,894 sites had greater asymmetry in PD patients (62.7%; *p* < 10^-15^, Fisher’s exact test; Fig. 2b). CpH sites were an important source of the hemispheric differences in DNA methylation (5,465 CpHs, 0.82 ± 0.02% change in PD; 742 CpGs, 4.40 ± 0.14% change in PD; *p* < 10^-26^ for CpH enrichment, hypergeometric test; Fig. 2c). In PD, hemispheric asymmetry in DNA methylation most often affected cis-acting regulatory elements, on average within 45.9 ± 1.0 kb of transcription start sites. We then identified the gene targets of the enhancers and promoters, using the Hi-C chromatin conformation data in prefrontal cortex and the *in silico* cis-regulatory element prediction tool, as described above. We found that PD patients had 5,728 genes with abnormal hemispheric asymmetry in epigenetic regulation, relative to controls (Supplementary Data 5). Moreover, DNA methylation abnormalities across PD hemispheres affected many PD risk genes^48^ (Fig. 2a). Thus, the neurons of PD patients exhibit extensive hemispheric asymmetry in the epigenetic regulation of genes, including known PD risk genes.

**Fig. 2.**
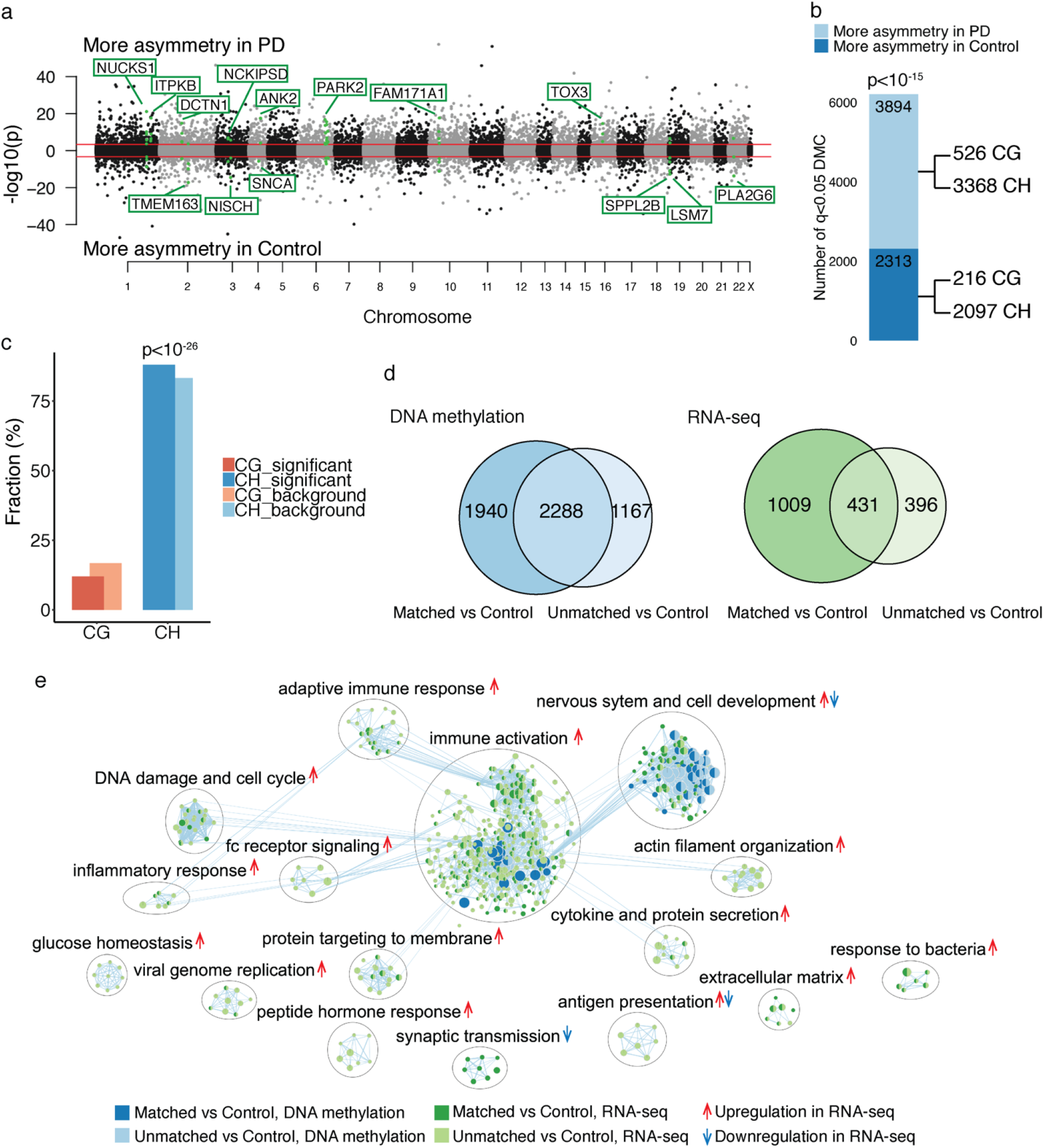
Epigenetic differences across hemispheres is increased in PD and is relevant to the lateralization of clinical symptoms. (**a**) Manhattan plot comparing the hemispheric asymmetry in DNA methylation in neurons of PD patients and controls, after controlling for age, sex, postmortem interval, and neuronal subtype proportion. DNA methylation differences were profiled at enhancers and promoters in prefrontal cortex neurons of 57 PD patients and 48 controls. −log10(p) refers to the significance of differentially methylated cytosines (DMC), with the sign corresponding to the diagnosis group with greater hemispheric asymmetry. Threshold for genome-wide significance (red line) is *q* < 0.05. Hemispheric differences in DNA methylation that affect genes linked to PD risk (familial and/or identified by GWAS^48^) are highlighted. (**b**) Comparison of the degree of hemispheric asymmetry in DNA methylation between PD patients and controls. The number of significant cytosine sites exhibiting more hemispheric asymmetry in PD or control neurons is shown. The *p*-value represents the enrichment of cytosines with greater hemispheric asymmetry in PD, by Fisher’s exact test. CpG and CpH contributions to hemispheric asymmetry are shown. (**c**) CpG and CpH involvement in hemispheric asymmetry changes in PD. The percent number of CpG or CpH sites significantly involved in hemispheric asymmetry changes in PD (relative to background) is shown. The enrichment of cytosine context was determined by hypergeometric test. (**d**) DNA methylation and transcriptional differences relevant to the lateralization of PD symptoms. Venn diagram showing the number of genes affected by differential methylation (left panel) or exhibiting differential expression (right panel) in the symptom-dominant (matched) or non-dominant (unmatched) PD hemisphere, relative to controls (DNA methylation: n = 17 PD-matched, 20 PD-unmatched, 48 controls; RNA-seq: n = 13 PD-matched, 11 PD-unmatched, 12 controls). (**e**) Pathway analysis of genes epigenetically and transcriptionally altered in neurons of the PD hemisphere matched or unmatched to side of symptom predominance, relative to control neurons. Pathway enrichment analysis of DNA methylation data was done by g:Profiler^92^ (blue nodes) and of RNA-seq data was done by GSEA preranked^93^ (green nodes). Epigenetically and transcriptionally dysregulated pathways in PD hemispheres were merged in Enrichment Map^94^ and annotated by Autoannotate^95^ using *q* < 0.05 pathways. Dark green or dark blue nodes show greater disruption in symptom-dominant PD hemisphere; light green or light blue nodes show greater disruption in symptom non-dominant PD hemisphere (relative to controls). Pathways up- or down-regulated in PD hemispheres depicted by red or blue arrows, respectively, as determined by GSEA^93^.

### Symptom lateralization involves epigenetic and transcriptional divergence between hemispheres

We examined whether DNA methylation abnormalities in PD were most apparent in the hemisphere matched to the side of clinical motor symptom predominance. Our highly characterized PD cases had information about the side of the body on which symptoms predominated (Supplementary Data 1). We profiled neurons of 17 PD patients with the hemisphere matched to the symptom-dominant side (the contralateral hemisphere) and 20 unmatched (the ipsilateral hemisphere), and we compared these PD groups to 48 control subjects. Neurons from the hemisphere matched to symptom dominance exhibited more DNA methylation differences relative to control hemispheres than did neurons from the unmatched hemisphere (Fig. 2d). There were 3,587 DNA methylation sites differing between the matched hemisphere of PD patients and controls, compared to 2,283 between the unmatched hemisphere of PD patients and controls (*q* < 0.05, robust linear regression with contrasts, adjusted for age, sex, postmortem interval, brain hemisphere (left or right), and neuronal subtype proportion; Supplementary Data 4). The identification of gene targets (as described above) revealed that differentially methylated regulatory elements in PD neurons affected 4,228 genes in the hemisphere matched to symptom predominance and 3,455 genes in the unmatched hemisphere, of which 2,288 genes were altered in both hemispheres (Fig. 2d; Supplementary Data 5). Overall, there were substantially more genes altered in the symptom-dominant hemisphere: 4,194 genes had more DNA methylation abnormalities at their regulatory elements in the PD hemisphere matched to symptom dominance than in the unmatched hemisphere (*q*<0.05 cytosines with fold change in matched *vs.* control > unmatched *vs.* control; Supplementary Data 5). Thus, prominent hemispheric asymmetry in the epigenomes of PD patient neurons mirrors the lateralization of clinical symptoms.

We determined whether epigenetic divergence between hemispheres is relevant to asymmetry in transcriptional patterns. We performed a transcriptomic analysis of the prefrontal cortex from control hemispheres and from PD hemispheres matched or unmatched to the symptom-dominant side (n = 12 controls, 13 PD-matched, 11 PD-unmatched; Fig. 2d). As in our epigenetic analysis, we found that the hemisphere matched to the symptom-dominant side had greater transcriptional differences relative to control hemispheres than did the unmatched hemisphere (Fig. 2d). Specifically, the matched PD hemisphere had 1,440 differentially expressed genes, while the unmatched PD hemisphere had 827 differentially expressed genes, compared to controls (*q* < 0.1, generalized linear regression with contrasts; Fig. 2d; Supplementary Data 6), after adjusting for age, sex, brain hemisphere, neuron proportion, RIN, and other sources of variation (by RUVSeq^49^). There were 1,121 genes with greater transcriptional changes in the matched PD hemisphere than in the unmatched hemisphere. There was also a significant association between changes in DNA methylation at gene regulatory elements and changes in corresponding transcript levels in PD (*p* < 0.05, interaction term in linear regression; Supplementary Fig. 3). Thus, hemispheric differences in the epigenome are accompanied by functionally relevant transcriptomic alterations.

We then determined convergent pathways affected by epigenetic and transcriptional changes involved in hemispheric asymmetry in PD. We identified pathways altered in PD hemispheres matched or unmatched to the side of symptom predominance relative to controls, and then compared these pathways between the PD hemispheres. In PD, there were inter-hemispheric differences in immune activation, inflammation, neuronal development, glucose homeostasis, protein localization, and synaptic activity (*q* < 0.05, g:Profiler and GSEA; Fig. 2e). Hence, asymmetry in PD may result from brain hemisphere differences in immune responses, neurodevelopment, and/or neurotransmission.

### Validation of hemispheric asymmetry in the epigenome with an independent PD and control cohort

To confirm our discovery that hemispheric asymmetry in DNA methylation in PD exceeds that of controls and corresponds to clinical motor symptom predominance, we replicated our findings with an independent cohort. In this replication study, we examined neurons from both the left and right hemisphere of the same person. DNA methylation was profiled in neuronal nuclei isolated from the prefrontal cortex of 31 controls and 26 PD patients: 12 PD patients with left-side symptom predominance and 14 with right-side predominance (Supplementary Data 1). Fine-mapping of DNA methylation at all brain enhancers and promoters was performed using the same bisulfite padlock probe approach as in our discovery cohort. After data preprocessing, DNA methylation was examined at 815,367 cytosine sites (133,736 CpGs and 681,631 CpHs; Supplementary Fig. 2). As above, we analyzed hemispheric asymmetry at individual cytosine sites (CpGs and CpHs) in neurons of the left and right hemisphere of PD patients and controls, adjusting for age, sex, postmortem interval, and neuronal subtype proportion.

In this independent cohort, we again found that there was more hemispheric asymmetry in neurons of PD patients than in controls (*q* < 0.05, robust linear regression with contrasts; Supplementary Fig 4b; Supplementary Data 4-5). DNA methylation abnormalities in PD also prevailed on the hemisphere side matched to symptom predominance (Fig. 3a). The symptom-dominant hemisphere had 589 differentially methylated cytosines, while the non-dominant hemisphere had 240 differentially methylated cytosines, relative to controls (*q* < 0.05, robust linear regression with contrasts; Supplementary Data 4). Genes targets of enhancers and promoters with hemispheric differences in DNA methylation in the replication cohort were identified as above. We found a total of 1,217 genes exhibiting more DNA methylation alterations in the symptom-dominant PD hemisphere (relative to controls) than in the non-dominant hemisphere (relative to controls) (Supplementary Fig. 4c; Supplementary Data 5). Moreover, there was a strong overlap between the discovery and replication cohort in the genes with DNA methylation abnormalities preferentially occurring in the symptom-dominant hemisphere (*p* < 10^-120^, hypergeometric test; Fig. 3b). This signifies an independent replication of epigenetically dysregulated genes involved in the lateralization of PD symptoms.

**Fig. 3.**
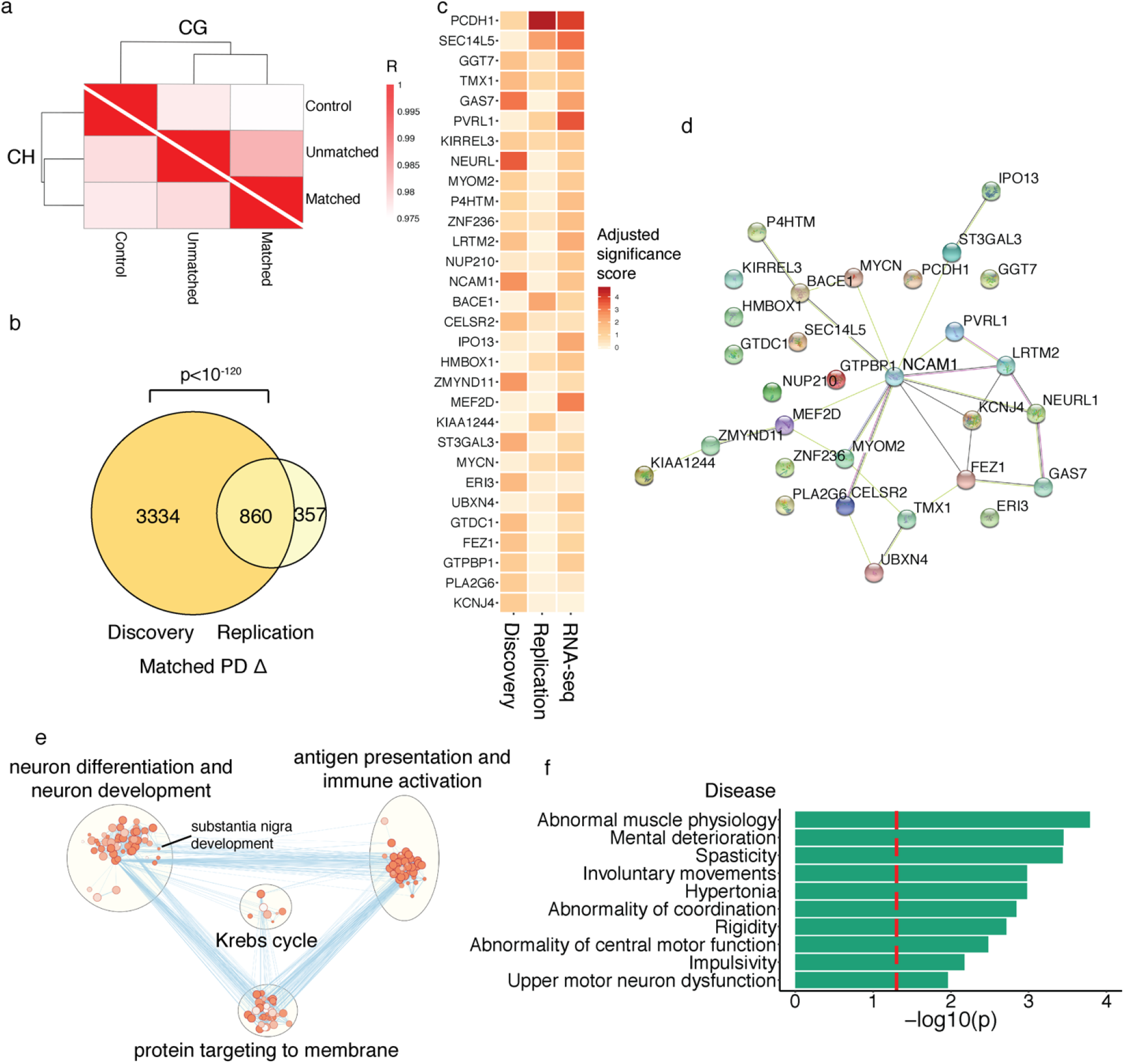
Independent validation that epigenetic dysregulation is greater in the PD hemisphere matched to side of symptom predominance. DNA methylation at enhancers and promoters was examined in an independent cohort of PD patients and controls. Prefrontal cortex neurons from both hemispheres were examined (n = 31 controls, 26 PD patients). (**a**) Divergence of epigenetic profiles in neurons of PD hemispheres matched or unmatched to the side of symptom dominance from those of control neurons. Shown are Pearson correlations comparing DNA methylation status of control, matched, and unmatched groups. Darker red signifies a higher DNA methylation similarity between diagnosis groups. (**b**) Concordance between the discovery and replication cohorts. Venn diagram showing the overlap of the discovery and replication cohorts for genes with greater epigenetic abnormalities in the symptom-dominant hemisphere of PD patients. The *p*-value represents the significance of the discovery and replication cohort overlap, by hypergeometric test. (**c**) Top 30 genes most associated with PD symptom lateralization. Genes were ranked across datasets to determine the most robustly dysregulated genes in the symptom-dominant hemisphere of PD patients. The discovery and replication DNA methylation data, as well as the RNA-seq data, was used. Genes are listed in ranked order, and the heatmap depicts the adjusted significance score in each dataset. (**d**) Network analysis of top 30 genes involved in symptomatic asymmetry in PD centers on *NCAM1*. Network analysis performed by STRING^96^. (**e**) Pathways of proteomic alterations involved in hemispheric asymmetry and symptom lateralization in PD. Protein changes were determined by mass spectrometry in the prefrontal cortex of PD patients, relative to controls, and between the PD symptom-dominant and non-dominant hemispheres (n = 3-5 individuals). Pathway enrichment analysis performed by g:Profiler^92^ (nodes are *q* < 0.05 pathways, hypergeometric test). Pathways were merged in Enrichment Map^94^ and annotated by AutoAnnotate^95^. (**f**) Top disease pathways of proteins involved in hemispheric asymmetry in PD. Pathway analysis was done by g:Profiler^92^. Threshold for significance (red dashed line) is *q* < 0.05.

**Fig. 4.**
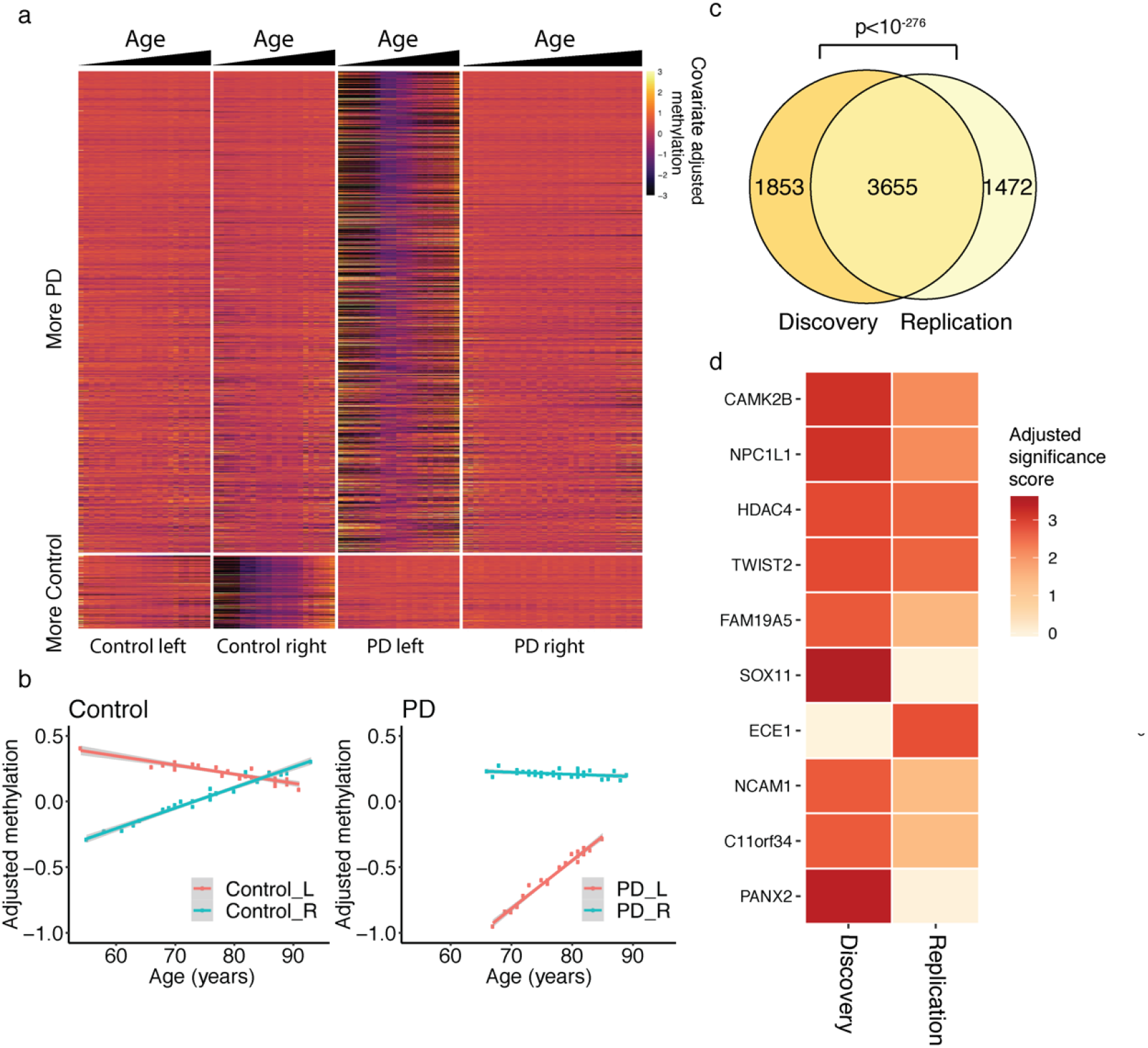
Progressive loss of hemispheric asymmetry in DNA methylation with aging. Age-dependent changes in DNA methylation in neurons of the left and right hemisphere of PD patients and controls (n = 23 PD-left, 34 PD-right, 25 control-left, 23 control-right). (**a**) Heatmap showing adjusted DNA methylation levels at the 5,925 cytosine sites exhibiting significant changes in hemispheric asymmetry with aging (*q* < 0.05, robust linear model followed by contrasts; DNA methylation adjusted for sex, postmortem interval and neuronal subtype proportion). Cytosine sites with more aging changes in hemispheric asymmetry in PD (upper panels) or in controls (lower panels) are shown. (**b**) Scatter plot of adjusted DNA methylation changes with age in the left and right hemispheres of controls and PD patients. The grey area represents confidence intervals. (**c**) Concordance in aging changes in DNA methylation in the discovery and replication cohort. Venn diagram showing the significance of overlap between the discovery and replication cohorts in the genes with age-dependent changes in hemispheric asymmetry in DNA methylation. The *p*-value represents significance of overlap by hypergeometric test. (**d**) Top 10 genes involved in aging changes in hemispheric asymmetry. Genes affected by aging changes in DNA methylation were ranked using the discovery and replication DNA methylation datasets. Genes are listed by ranked order, and the heatmap shows adjusted rank significance score in each dataset.

We sought to identify the genes that were most strongly associated with asymmetry in PD symptoms. We used the epigenetic and transcriptomic data from our discovery cohort and the epigenetic data from our replication cohort, and searched for the genes most dysregulated in the hemisphere corresponding to the side of symptom dominance. Each gene was ranked according to the significance of change and consistency across the epigenetic and transcriptomic datasets. We identified 49 genes showing consistent preferential dysregulation in the hemisphere matched to symptom predominance (Fig. 3c and 3d; Supplementary Data 7). In particular, the neuronal cell adhesion molecule 1 (*NCAM1*), which regulates neuronal development, synaptogenesis, cell–cell interactions, and synaptic plasticity^50^, was central to the abnormalities in the symptom-dominant PD hemisphere (Fig. 3d).

### Proteomic analysis of hemispheric asymmetry in PD

To further explore the genes involved in hemispheric asymmetry in PD, we performed a quantitative proteomic analysis of the PD prefrontal cortex. We first identified 1,063 proteins with altered abundances in the prefrontal cortex of PD patients relative to controls (n = 3 PD patients and 3 controls; Supplementary Data 8). We then determined 668 protein differences between the PD hemispheres matched and unmatched to symptom dominance (n = 5 PD-matched and 5 PD-unmatched). These analyses were merged to identify 345 disease-relevant proteins that exhibit hemispheric asymmetry in the PD brain (Supplementary Data 8). Notably, we identified that SNCA (α-synuclein) and NCAM1 were altered in PD and exhibited hemispheric asymmetry (Supplementary Fig. 5). SNCA and NCAM1 levels were highest in the PD hemisphere matched to symptom dominance (Supplementary Fig. 5; Supplementary Data 8). Pathway analysis of the PD-relevant proteins exhibiting hemispheric asymmetry revealed differences in nervous system development (including substantia nigra development), antigen presentation and immune activation, and protein transport (*q* < 0.05, hypergeometric test; Fig. 3e). Moreover, disease pathways associated with the protein changes involved in hemispheric asymmetry in PD were related to motor dysfunction, including many hallmark symptoms of PD (9 of the top 10 human disease pathways involved in motor dysfunction; *q* < 0.05, hypergeometric test; Fig. 3f). Therefore, proteomic analysis supports the epigenetic and transcriptomic findings that symptom asymmetry in PD is associated with differences in neurodevelopmental processes and immune responses between hemispheres.

### Changes in hemispheric asymmetry in healthy and PD neurons with aging

In PD, highly lateralized motor symptoms gradually become more bilateral with increased age and disease duration; though lateralization persists even in advanced PD stages^10, 11^. We found that hemispheric asymmetry in the epigenome of prefrontal cortex neurons changed with aging, especially for PD patients (Fig. 4). We found 5,925 methylated cytosines at enhancers that showed changes in hemispheric asymmetry with aging (*q* < 0.05, robust linear regression with contrasts; adjusted for sex, postmortem interval, and neuronal subtype proportion; n = 23 PD-left, 34 PD-right, 25 control-left, 23 control-right; Fig. 4a; Supplementary Data 4). When these sites were categorized according to diagnosis and hemisphere side, we observed that there are more aging changes in hemispheric asymmetry in the epigenomes of PD patients (5,146 out of 5,925 significant sites showing greater aging changes in PD than in controls; *p* <10^-15^, Fisher’s exact test; Fig. 4a). The left hemisphere of PD patients was particularly vulnerable to aging changes in DNA methylation, concordant with previous PD imaging studies^5,51–54^. Interestingly, in both PD patients and controls, DNA methylation at enhancers and promoters became increasingly symmetrical across hemispheres with aging (Fig. 4b; Supplementary Fig. 6). There was a reduction in asymmetry with aging at 4,198 (70.9%) cytosines in PD patients and 3,818 (64.4%) cytosines in controls, out of the 5,925 age-associated sites (*p* < 0.001 and *p* < 10^-7^, respectively, Fisher’s exact test; Fig. 4b). Though neuronal epigenomes across PD hemispheres became less asymmetric with age, left–right asymmetry in PD neuronal epigenomes persists even at advanced ages (Fig. 4b). Therefore, in aging, there is a progressive loss of hemispheric asymmetry in neuronal epigenomes in both controls and PD patients. Loss of epigenetic asymmetry between hemispheres of PD patients may contribute to bilateral symptomatic progression in PD^11^.

We then identified the genes that had the most robust aging changes in hemispheric asymmetry in PD. We first determined the gene targets of the regulatory elements showing hemispheric differences in DNA methylation with aging and PD diagnosis (using the chromatin conformation analysis approach described above). There was strong overlap in the discovery and replication cohorts in the genes that showed aging changes in hemispheric asymmetry in PD relative to controls (*p* < 10^-276^, hypergeometric test; Fig. 4c; Supplementary Data 5). The genes with the most significant aging changes in hemispheric asymmetry in PD were then ranked based on consistency across the discovery and replication cohorts. Among the top genes, there were calcium/calmodulin-dependent protein kinase 2 (*CAMK2B*), histone deacetylase 4 (*HDAC4*), and *NCAM1*, which have established roles in synaptic plasticity, neurotransmitter release, neurodevelopment, memory, and locomotor activity^50, 55–57^, as well as endothelin-converting enzyme-1 (*ECE1*), which degrades α-synuclein pathology^58^ (Fig. 4d). Hence, PD patients have aging changes in the hemispheric asymmetry of DNA methylation that affect genes involved in synaptic transmission, motor functions, and α-synuclein levels.

### Differential hemispheric asymmetry in the epigenome is associated with PD progression

We examined the relationship between hemispheric asymmetry and disease course in PD patients. A short disease course was defined as less than 15 years of PD motor symptoms prior to death, while a long disease course exceeded 15 years. PD patients with either a short or a long disease course had a similar ages at death (average age, short course: 77.4 ± 1.7 years; long course: 77.3 ± 1.7 years; both hemispheres of n = 14 and 12 PD patients with short or long disease course, respectively; Supplementary Data 1). In PD neurons, we examined the divergence of DNA methylation patterns between the symptom-dominant and non-dominant hemispheres with PD disease course. There were 2,910 cytosine sites in enhancers and promoters showing changes in hemispheric asymmetry with PD duration (*q* < 0.05, robust linear regression with contrasts, adjusting for age, sex, postmortem interval, brain hemisphere (left or right), and neuronal subtype proportion; Supplementary Data 4). The hemispheric asymmetry in DNA methylation was greater in neurons of PD patients that had a long disease course (*p* < 0.001, Student’s *t*-test; Fig. 5a). Hence, prominent epigenetic differences between hemispheres are linked to a slow PD progression. The greater hemispheric asymmetry in the epigenomes of PD patients with a longer disease course may explain the clinical observations that PD patients with highly lateralized symptoms have a slower disease progression than those with symmetrical symptoms^12^.

**Fig. 5.**
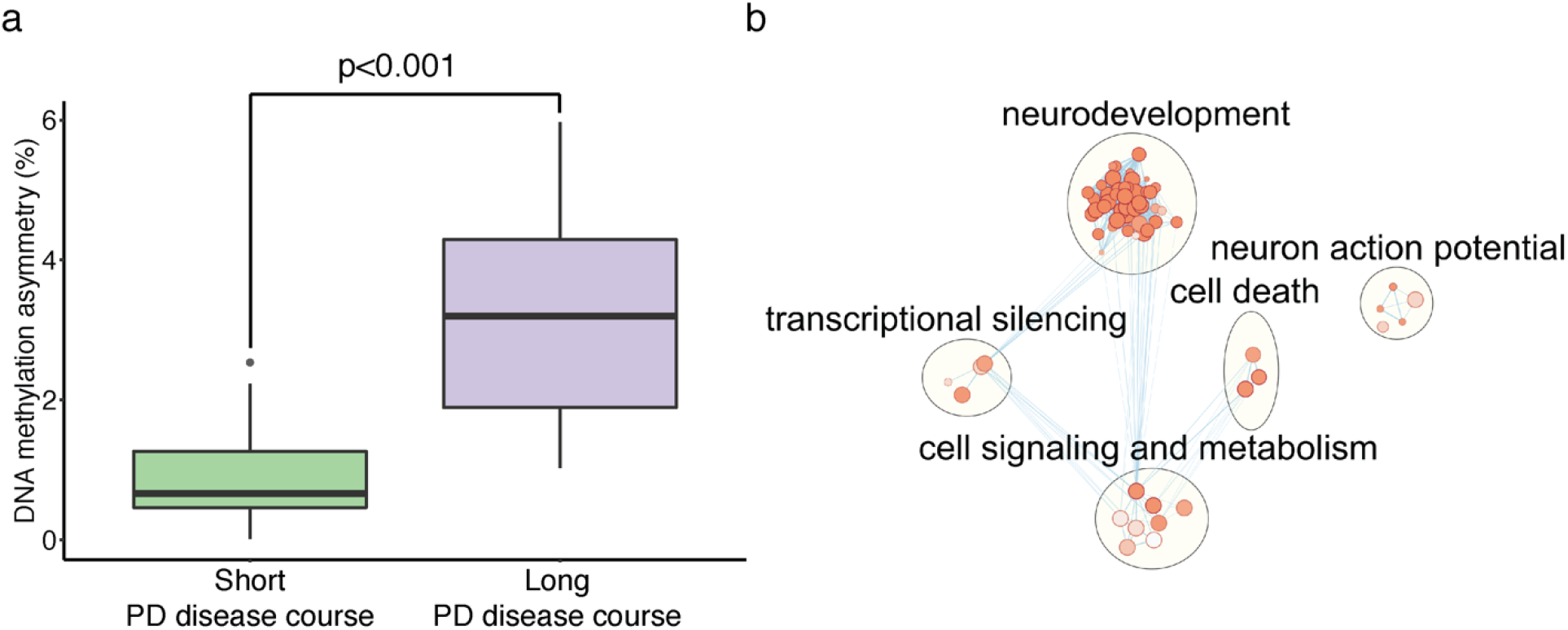
A long PD disease course is associated with high levels of hemispheric asymmetry in DNA methylation. (**a**) The extent of hemispheric asymmetry in DNA methylation in PD patients with a short or long disease course. Significant DNA methylation changes associated with PD duration were identified (n = 2,910 cytosine sites, *q* < 0.05, robust linear regression with contrasts, controlling for age, sex, postmortem interval, neuronal subtype proportion, and brain hemisphere side). The extent of hemispheric asymmetry for cytosine sites associated with PD duration was determined in PD patients with a short (≤15 years) or long (>15 years) disease course (n = 13 and 11 individuals, respectively). The boxplot center line is the median, the lower and upper limits are the first and third quartiles (25^th^ and 75^th^ percentiles), and the whiskers are 1.5 × the interquartile range. *p* < 0.001 is the difference between the short and long disease course groups in level of DNA methylation asymmetry, as determined by *t*-test. (**b**) Pathways differing between PD patients with a short or long disease course. Pathway enrichment analysis of genes with epigenetic differences associated with PD duration was performed by g:Profiler^92^ (nodes are *q*<0.05 pathways, hypergeometric test). Pathways were merged in Enrichment Map^94^ and annotated by AutoAnnotate^95^.

Genes and pathways in neurons associated with differences in PD disease course were investigated. The gene targets of enhancers and promoters that showed hemispheric asymmetry changes in DNA methylation with PD disease course were identified (using Hi-C analysis of prefrontal cortex and the *in silico* regulatory element prediction tool). There were 3,953 genes that had inter-hemispheric changes in epigenetic regulation with PD duration (Supplementary Data 5). Pathway analysis determined that the length of disease course was related to epigenetic changes at genes affecting neurodevelopment, neurotransmission, apoptosis, cell signaling, and metabolism (*q* < 0.05, hypergeometric test; Fig. 5b). Hence, epigenetic mechanisms impacting brain development and neuronal communication may influence the progression of PD.

### PD risk genes exhibit hemispheric asymmetry in DNA methylation

Finally, we sought to understand the contribution of PD risk genes to hemispheric asymmetry in PD. PD risk genes (determined by GWAS meta-analysis^48^) were identified among the genes that exhibited greater hemispheric asymmetry in PD, relative to controls, and that were preferentially disrupted in the symptom-dominant PD hemisphere. We also examined the contribution of PD risk genes to aging-related and disease-duration-related changes in hemispheric asymmetry in PD patients. Our DNA methylation, transcriptomic, and proteomic analyses were used to identify hemispheric differences in PD patients that involved PD risk genes (significant genes in analysis for Fig. 2-5; Supplementary Data 5). We found 34 out of 72 PD risk genes showing more hemispheric asymmetry in PD relative to controls, and/or showing greater differences in the PD hemisphere matched to symptom dominance (Fig. 6). These included genes involved in immune cell functioning and development (*ITPKB, SATB1*)^59, 60^, axonal growth and synaptic signaling (*ANK2*, *CAMK2D*)^61, 62^, and α-synuclein pathology (*SNCA*). Similar PD GWAS risk genes exhibited inter-hemispheric methylation changes with aging and were linked to a short duration of PD (Fig. 6). We also noticed that a region on chromosome 3 (∼300 kb, spanning from *ALAS1* to *STAB1*) was consistently involved in hemispheric asymmetry in PD across datasets, suggesting a combined epigenetic and genetic disruption of this area in PD. These findings suggest that epigenetic changes at PD risk genes contribute to lateralized hemisphere dysregulation in PD.

**Fig. 6.**
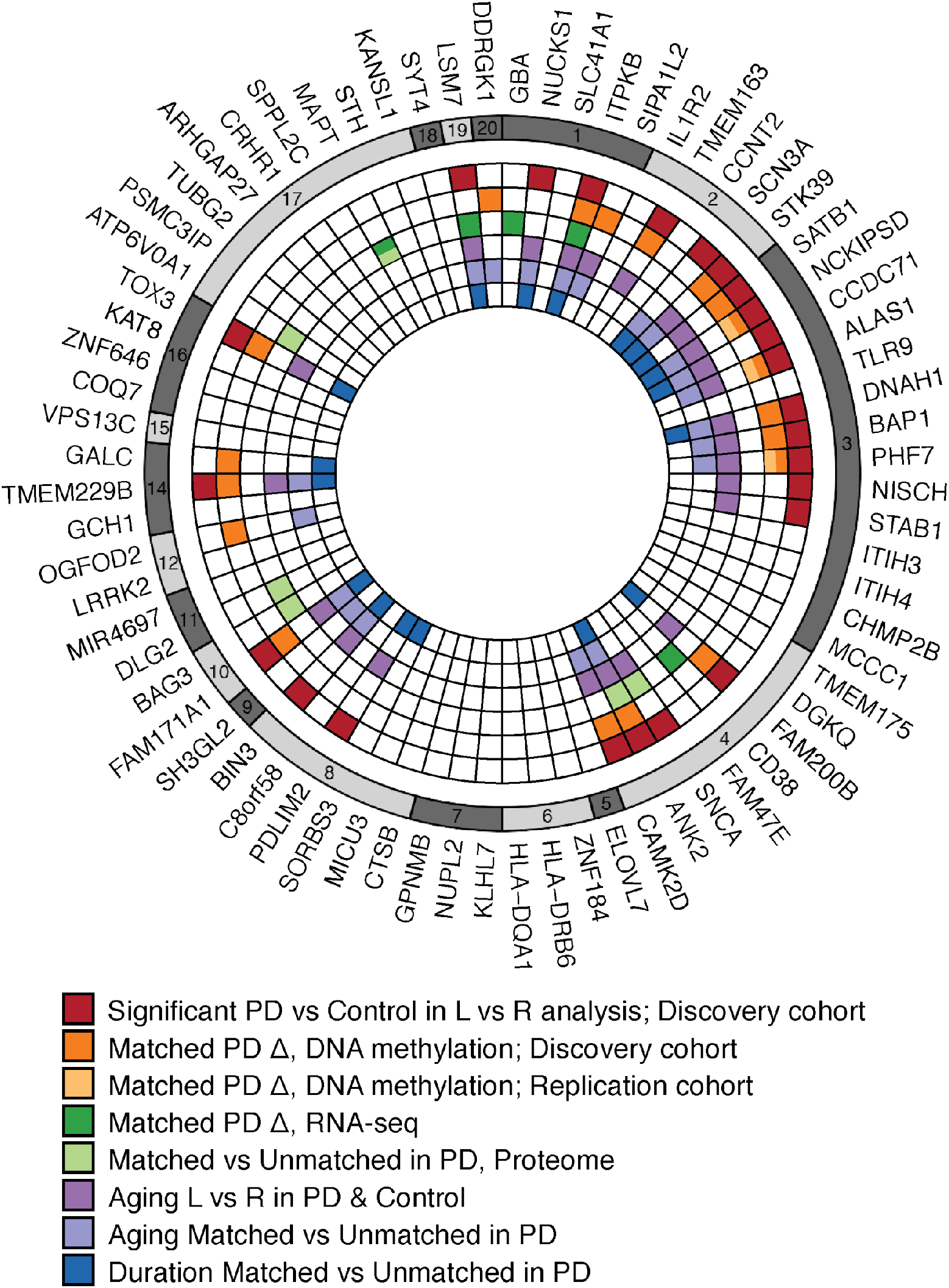
PD risk genes contribute to hemispheric asymmetry in neurons of PD patients. PD risk genes identified by GWAS^48^ were found to exhibit differential hemispheric asymmetry in PD patients relative to controls, were preferentially dysregulated in the PD symptom-dominant hemisphere, and were involved in hemispheric asymmetry changes occurring with aging and PD disease course. Genes identified in eight independent analyses that include epigenetic, transcriptomic, or proteomic data are presented. The plot summarizes PD risk genes with: 1) left–right hemisphere differences in PD relative to controls that involve epigenetic changes at cis-regulatory elements (red); 2 and 3) epigenetic dysregulation that is greater in the symptom-dominant hemisphere than in the non-dominant hemisphere (relative to control hemispheres), as determined in discovery cohort (orange) and replication cohort (yellow); 4 and 5) disruption in PD associated with symptom lateralization, as determined by analysis of the transcriptome (dark green) and proteome (light green); 6) aging changes in left–right hemispheric asymmetry in epigenetic regulation in PD and controls (purple); (7) aging changes in epigenetic regulation between the symptom-dominant and non-dominant hemisphere in PD (indigo); (8) hemisphere asymmetry in DNA methylation associated with length of PD disease course (blue). Colored boxes represent significant changes affecting PD risk gene.

We also determined the effects of cis-acting genetic variation on the DNA methylation changes involving hemispheric asymmetry in PD. SNPs that were proximal to the cytosine sites profiled in our bisulfite sequencing data were identified; 69% of cytosines related to hemispheric asymmetry in PD, and 70% of all cytosines profiled in our study, had one or more SNPs identified within ± 500 kb. We then performed a methylation quantitative trait loci (me-QTL) analysis to examine the effects of genotype on DNA methylation and determined that 11,507 out of 564,294 cytosines had a meQTL association (*q*<0.05, robust linear regression, after adjusting for diagnosis, hemisphere, age, sex, postmortem interval, and neuronal subtypes). We found that cytosine sites relevant to hemispheric asymmetry in PD had a significant increase in SNP associations relative to all cytosine sites profiled (4.5% of cytosines related to hemispheric asymmetry in PD *vs.* 2.0% of background cytosines, *p* < 10^-15^, χ^2^ test). Thus, genetic factors may drive a number of DNA methylation changes involved in hemispheric asymmetry in PD.

## Discussion

Although hemispheric asymmetry is a fundamental biological feature of the human brain, its determinants remain unclear. Here, we demonstrate that the epigenetic states of neurons differ between hemispheres and may play a role in the lateralization of brain functions. The prominent inter-hemispheric differences in the epigenome, transcriptome, and proteome identified in this study signify that hemisphere side should be considered in future molecular studies of brain health and disease.

Our analysis showed hemispheric differences in DNA methylation in neurons of the healthy prefrontal cortex were mainly driven by differential accumulation of CpH methylation. At enhancers, CpH methylation acts as an epigenetic silencer and is highly associated with gene transcript levels^21, 23^. The accumulation of CpH methylation, which we found to preferentially occur in neurons of the left hemisphere, affected genes involved in neurodevelopment, synaptic organization, and neurodegeneration. Neuronal asymmetry in CpH silencing of gene activity may instill a differential vulnerability of hemispheres to synaptic dysfunction and neuronal loss that lead to neurodegenerative diseases. In support, hemispheric asymmetry in PD was largely driven by changes in CpH methylation.

Our study shows that hemispheric asymmetry of the epigenome is exaggerated in PD, with greater methylation abnormalities observed in the hemisphere matched to the symptom-dominant side. A major challenge in epigenetic studies is differentiating epigenetic changes that are causal to disease from those that arise as a consequence of a non-shared environment. Because we examined neurons in each brain hemisphere of the same individual, we were able to delineate epigenetic differences that preferentially appear on the symptom-dominant side of the PD brain from those appearing in both hemispheres. As such, we identified epigenetic changes, genes, and pathways most relevant to the presentation of PD clinical symptoms. We reinforced our findings with transcriptomic and proteomic analyses and replicated our results in an independent cohort. Finally, hemispheric asymmetry in the epigenome impacts PD risk genes identified by GWAS^48^, further supporting that epigenetic dysregulation at these genes could contribute to disease pathobiology.

In advanced age, there is a convergence of epigenetic changes between hemispheres in healthy and PD neurons. This could explain the reduction in hemispheric asymmetry observed in functional neuroimaging studies of old as compared to young adults^63, 64^. We also found that the left hemisphere of the PD brain had greater epigenetic dysregulation than the right hemisphere, which is consistent with PD imaging studies^63, 64^. The convergence of hemisphere epigenomes in aging PD patients may explain why symptoms become more bilateral as PD progresses^11^. We also found that a long PD duration was associated with greater hemispheric asymmetry in DNA methylation. Clinically, PD patients with symmetrical symptom onset are prone to rapid disease progression^12^ and, in our study, rapid disease progression was associated with an epigenetic disruption of genes involved in neurodevelopment, cell signaling, and cell death.

Based on the results of our multiomics study, we postulate that hemispheric asymmetry in PD results from the differential regulation of genes involved in nervous system development, immune signaling, and synaptic transmission (Fig. 7). Lateralization is developed early in the brain, and neuronal progenitors can shape the recruitment and positioning of brain-resident immune cells, the microglia^65, 66^. These immune cells are sessile, show regional differences in the brain^67, 68^ and have dynamic processes that shape synaptic structure, maturation, and signaling^69, 70^. Hence, it is possible that early established differences in neuronal structure, in combination with lifelong differences in immune activity and neurotransmission across hemispheres, leads to unilateral vulnerability, which may explain the corresponding unilateral presentation of PD symptomatology.

**Fig. 7.**
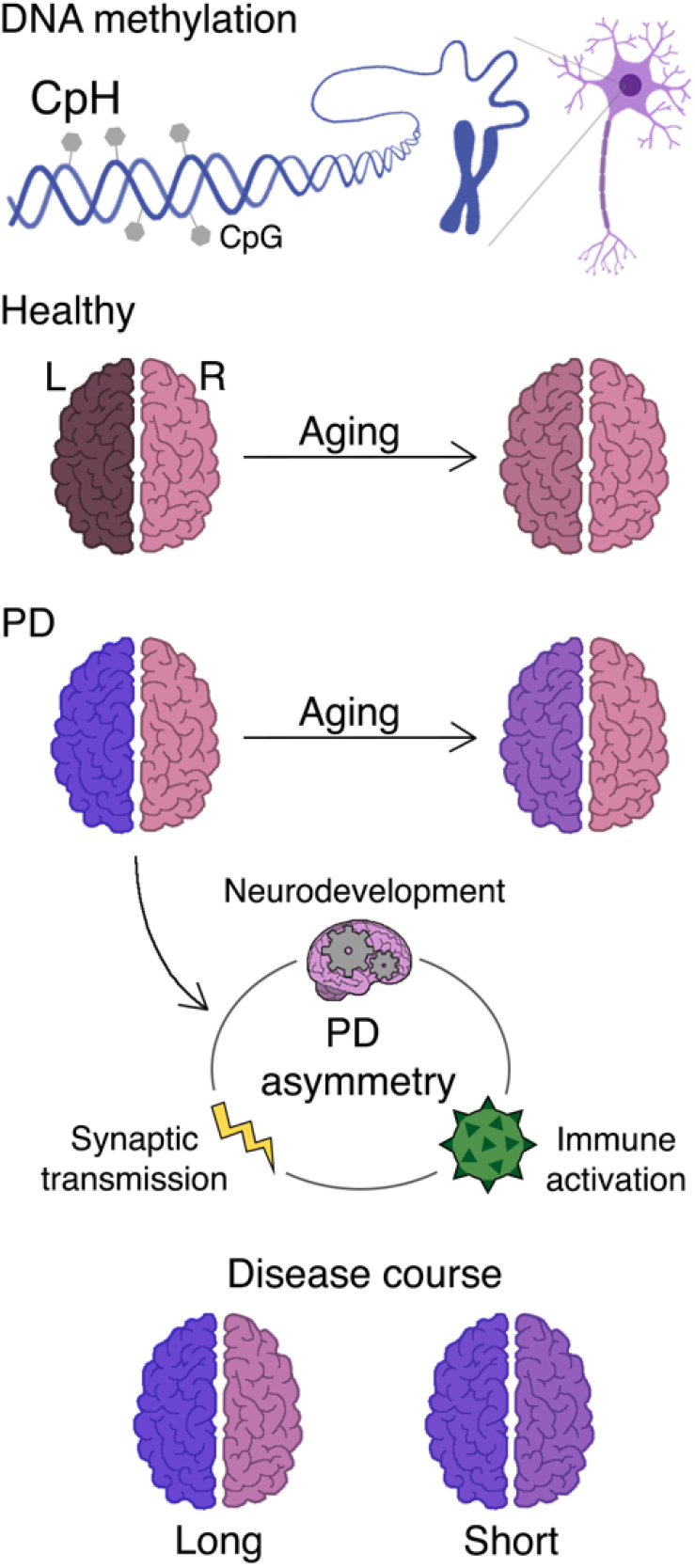
Schema of hemispheric asymmetry in the epigenome of the healthy and PD brain. Neurons of the healthy human brain exhibit prominent hemispheric asymmetry in DNA methylation. The healthy brain has higher DNA methylation levels in the left hemisphere, a difference driven largely by the accumulation of CpH methylation. Compared to neurons in the healthy brain, neurons in the PD brain possess considerably greater hemispheric asymmetry, again primarily driven by differential CpH methylation. Hemispheric asymmetry in PD involves DNA methylation abnormalities that are more prominent on the hemisphere matched to the side of symptom dominance. Aberrant hemispheric asymmetry and symptom lateralization in PD is related to disruption of genes affecting neurodevelopment, immune activation, and synaptic transmission. In aging, neuronal epigenomes exhibit a decrease in hemispheric asymmetry. The convergence of neuronal epigenomes in PD with aging may contribute to the bilateralization of PD symptoms over time, though hemispheric asymmetry in DNA methylation persists even at advanced ages. Epigenetic asymmetry between hemispheres is also linked to disease progression: PD patients with long (>15 years) disease courses have greater asymmetry than patients with short (≤15 years) disease courses. Shading of brain hemispheres represents asymmetry in DNA methylation between paired hemispheres (shading represents DNA methylation status).

## Acknowledgments

We thank the Van Andel Research Institute Flow Cytometry, Genomics, Bioinformatics and Biostatistics, and Pathology Cores. We thank Zachary Madaj in the Bioinformatics and Biostatistics Core at the Van Andel Research Institute for consulting on statistical approaches. We also thank the CAMH Sequencing Facility. We appreciate the manuscript reviews provided Dr. Art Petronis and Edward Oh. We thank the patient donors and families for the support of our research and the Parkinson’s UK Brain Bank, the NIH NeuroBioBank, and the Michigan Brain Bank for the brain tissue provided. V.L. is supported by grants from the Department of Defense (W81XWH1810512), the National Institute of Neurological Disorders and Stroke (1R21NS112614-01), and a Gibby & Friends vs. Parky Award.

## Data Availability

All sequencing data generated in this study are available from the NCBI Gene Expression Omnibus (GEO) database under the accession numbers GSE135331 and GSE135037. The mass spectrometry proteomics data have been deposited to the ProteomeXchange Consortium via the PRIDE^71^ partner repository with the dataset identifiers PXD015079 and PXD015239. Custom code for DNA methylation and RNA-seq analysis is available at https://github.com/lipeipei0611/PD_Asymmetry.

## Author contributions

PL contributed to experimental design and performed all computational analyses. EE performed the neuronal nuclei isolations and contributed to bisulfite padlock probe library preparation. LM contributed to the RNA-seq analysis. MS, JL, and IV performed the proteomics analysis. VL was involved with study design and overseeing the experiments. The manuscript was written by VL, SL, and PL, and all authors commented on the manuscript.

## Online Methods

No statistical methods were used to predetermine sample size.

### Human tissue samples

Human prefrontal cortex tissue for this study was obtained from the Parkinson’s UK Brain Bank, NIH NeuroBioBank, and Michigan Brain Bank, with approval from the ethics committee of the Van Andel Research Institute (IRB #15025). For each individual, we had information on demographics (age, sex, ethnicity), brain hemisphere, tissue quality (post-mortem interval), side of symptom predominance, PD duration, and pathological staging (Supplementary Data 1). Control individuals had pathologically normal brains (and had no brain Lewy body pathology). PD cases were pathologically confirmed to have brain Lewy body pathology. We examined two independent cohorts of samples; the discovery and replication cohort. The discovery cohort included 105 individuals: 48 controls (brain hemisphere left: 25; right: 23) and 57 PD patients (brain hemisphere left: 23; right: 34). For the PD patients in the discovery cohort, 37 had information about the side of symptom predominance (hemisphere matched to side of symptom dominance: 17; unmatched: 20). The replication cohort included both hemispheres of 57 individuals: 31 controls and 26 PD patients, and prefrontal cortex tissue was obtained from the same area of both hemispheres for each individual. The side of symptom predominance was known for all the PD patients in the replication cohort. Neurons of the prefrontal cortex were selected for this study because 1) this brain region plays an important role in PD^72^ 2) pathology does spread to this brain region^73^, and 3) neurons remain present in the PD prefrontal cortex^74^ (in contrast to the substantia nigra, which has undergone severe neurodegeneration^11^).

### Isolation of neuronal nuclei by flow cytometry

Neuronal nuclei were isolated from the human prefrontal cortex using flow cytometry^32, 45^. Fresh-frozen prefrontal cortex tissue (200-300 mg) was rinsed and finely chopped in 2 mL of PBSTA Buffer (0.3 M sucrose, 1× Dulbecco’s PBS (Gibco), 3 mM MgCl_2_). The sample was then homogenized on ice for three intervals of 5 s (BioSpec Tissue Tearor, on lowest setting). For each sample, 40 µL of 10% Triton X-100 was added to the homogenate and incubated for 15 min. Next, the tissue homogenate was transferred to a dounce homogenizer (Kimble) and homogenized 8 times. The homogenate was filtered through Miracloth (Calbiochem) and passed through a sucrose cushion (1.4 M, 1× Dulbecco’s PBS (Gibco), 0.1% TritonX-100, 3 mM MgCl_2_) by centrifugation at 3000 × *g* for 30 min at 4°C. After removing the supernatant, the pelleted nuclei were incubated for 15 min in 800 µL of blocking buffer: 1× PBS, 1.25% goat serum (Gibco), 3 mM MgCl_2_, and 0.0625% BSA (Thermo Fisher Scientific). The nuclei were then gently resuspended and mixed with anti-NeuN antibody (1:500, Abcam) and incubated for at least 30 min on ice. Immediately before sorting, 10 µL of 7-AAD or DAPI (Thermo Fisher Scientific, Sigma-Aldrich) was added to each sample, and nuclei samples were filtered through a 41-µm filter (Elko Filtering Co.). Samples were sorted on a MoFlo Astrios in the Flow Cytometry Core of the Van Andel Research Institute, using the gating strategy described in Supplementary Fig. 1. After sorting, nuclei were pelleted in 10 mL Dulbecco’s 1× PBS (Gibco), 0.3 M sucrose, 5 mM CaCl_2_, and 3 mM MgCl_2_. Samples were mixed by inverting and incubated on ice for 15 min before centrifuging at 2500 × *g* for 10 min. The supernatant was removed, and pellets were frozen at −80°C until DNA isolation. Neuronal nuclei DNA was isolated using standard phenol–chloroform methods.

### Fine-mapping of DNA methylation with bisulfite padlock probe sequencing

The bisulfite padlock probe sequencing technique was used for the targeted quantification of DNA methylation with single-nucleotide resolution at enhancers and promoters in neurons of the human prefrontal cortex^32, 75^. Human brain enhancers and promoters were identified using the EpiCompare tool^76^, which identifies tissue/cell type gene regulatory elements based on chromatin state data defined by the ChromHMM tool^77^ from the RoadMap Epigenomics Project^78^. Enhancers and promoters were defined based on the 18-state ChromHMM model^77^. Our study included all genic, active, weak, or poised/bivalent enhancers (7_EnhG1, 8_EnhG2, 9_EnhA1, 10_EnhA2, 11_EnhWk, 15_EnhBiv). We also included all promoters that were active, near a transcription start site, or poised/bivalent (1_TssA, 2_TssFlnk, 3_TssFlnkU, 4_TssFlnkD, 14_TssBiv; for E073, E072, and E074). Enhancers and promoters significantly enriched in the adult brain are from the Tissue Specific Enhancers website (https://epigenome.wustl.edu/TSE/browse.php). We also included all enhancers and promoters present in adult prefrontal cortex (E073), inferior temporal lobe (E072), and substantia nigra (E074).

Padlock probes (n = 59,009) for bisulfite analysis targeted the unique (non-repetitive) enhancer and promoter regions on both forward and reverse DNA strands. Padlock probes were designed using ppDesigner^79^ (v2.0) with the human GRCh37/hg19 genome. Probe sequences are described in Supplementary Data 2. Padlock probes were synthesized using a programmable microfluidic microarray platform (CustomArray, Inc.) and were prepared and purified for experiments, as described^75^.

DNA methylation fine-mapping using the bisulfite padlock probe sequencing approach was performed as previously described^32, 75^. In brief, genomic DNA for each sample was bisulfite-converted and purified using the EZ DNA Methylation Kit (Zymo Research). The bisulfite-converted DNA (200 ng) was hybridized to the padlock probes (1.5 ng). Targeted regions were extended using PfuTurbo Cx (Agilent Technologies), and circularization was completed using Ampligase (Epicentre). Non-circularized DNA was digested using an exonuclease cocktail, and the remaining target circularized DNA was amplified using a common linker sequence in the padlock probe. Libraries were PCR-amplified, purified with AMPure XP beads (Beckman Coulter A63881), pooled in equimolar amounts, and further purified using a QIAquick Gel Extraction kit (Qiagen). Libraries were quantified using the Qubit dsDNA HS Assay Kit (Thermo Scientific) and qPCR (Kapa Biosystems) on a ViiA 7 Real-time PCR system (Applied Biosystems). Next-generation sequencing of the libraries was performed by the Epigenetics Lab at the Centre for Addiction and Mental Health in Toronto, Canada, on an Illumina HiSeq 2500 machine in HiOutput mode. Library sequencing was done across 3 flow cells (24 lanes) for the discovery cohort and across 2 flow cells (16 lanes) for the replication cohort, yielding 25-40 million reads/sample.

### Epigenomic data analysis

We examined DNA methylation status at every cytosine site (CpG and CpH) covered by padlock probes targeting 35,288 regulatory regions across the genome with a custom pipeline^32, 75^. This pipeline was used for both the discovery (n = 108 unique samples, 17 technical replicates) and replication (n = 114 unique samples, 12 technical replicates) cohorts. First, we removed low-quality bases and performed adapter trimming of bisulfite-treated sequencing reads using Trimmomatic-0.32 for the discovery cohort or Trim Galore (v0.4.4) for the replication cohort. Bismark^80^ (v0.17.0) was used to align reads to the target reference genome (GRCh37/hg19) and perform methylation calls. Methylation calls were included only for cytosines with a minimum read depth of 30×. Bisulfite conversion efficiency was 99.14 ± 0.005% in the discovery cohort and 99.28 ± 0.007% in the replication cohort (averaged CC methylation per sample). We excluded 5 samples in the discovery cohort (3 unique samples and 2 replicates) and 3 samples in the replication cohort (3 unique samples) from further analyses due to poor inter-sample correlations (> 10% difference). Technical replicates confirmed a high reproducibility in the sample-level DNA methylation correlation analysis (average R for the discovery cohort: 0.94 ± 0.007; average R for the replication cohort: 0.97 ± 0.003; Supplementary Fig. 2). DNA methylation calls at each site were merged for matched technical replicate samples. Cytosine sites with missing (unknown) DNA methylation calls in more than 30% of samples were excluded. We also removed cytosine sites overlapping common SNPs (minor allele frequency ≥ 0.05), as identified by the 1000 Genomes Project (phase 3 v5a 20130502 release for chr1∼chr22, v1b 20130502 for chrX; all populations and European populations)^81^. CpG and CpH sites that had stable DNA methylation calls (DNA methylation status of 0% or 100% or NA) in more than 50% of samples were excluded from further analysis. At the end of these preprocessing steps, the discovery cohort had 105 samples with 633,803 CpGs/CpHs and the replication cohort had 111 samples with 815,367 CpGs/CpHs that consisted of quality-controlled, genome-wide methylation data retained for downstream analysis (Supplementary Data 1; Supplementary Fig. 2).

### Neuronal subtype proportion

In the prefrontal cortex, 70-85% of neurons are excitatory glutamatergic neurons, while the remaining 15-30% are inhibitory GABAergic neurons^82^. Our DNA methylation analysis adjusts for variation in neuronal subtypes. We performed cell-type deconvolution using CIBERSORT^83^ (http://cibersort.stanford.edu) and reference neuronal subtype-specific markers (gene body CpH methylation) provided in a single-cell DNA methylome analysis of the human frontal cortex^82^. For the reported 1,012 neuronal subtype gene signatures, we averaged CpH methylation within gene bodies (± 100 kb) and found 564 neuronal subtype gene signatures for the discovery cohort and 610 signatures for the replication cohort. Using the neuronal subtype signature matrix (gene CpH markers), CIBERSORT was run with 100 permutations. We did not find any significant differences in any type of neuron (Supplementary Data 3). To control for neuronal subtype variation in our DNA methylation analysis, we used the proportion of glutamatergic relative to GABAergic subtypes for each sample.

### Statistical analysis for differentially methylated sites

DNA methylation analysis for the discovery and replication cohort involved multivariate robust linear regression models with empirical Bayes from the limma (v3.30.13) statistical package^84^. DNA methylation was transformed from B-values to M-values using lumi (v2.30.0)^85^. Because the replication cohort data has both hemispheres from the same individual, we added in the limma model a blocking factor to define samples from the same individual and a correlation coefficient determined by the duplicateCorrelation function. For each dataset and contrast, *p* values were adjusted with a Benjamini-Hochberg correction for multiple testing and those with FDR *q* < 0.05 were deemed significant.

### Analysis of hemispheric asymmetry in DNA methylation

#### In controls, PD, and aging (model 1)

To analyze hemispheric asymmetry in DNA methylation, the linear model matrix for CpG/CpH methylation (M) as a dependent variable is:

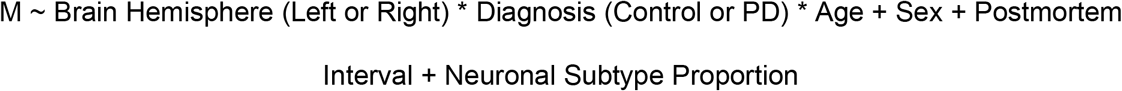

We used the contrasts.fit function to identify sites exhibiting hemispheric asymmetry in controls with the following contrasts matrix: Right brain hemisphere in controls **–** Left brain hemisphere in controls. To identify sites exhibiting hemispheric asymmetry in PD patients relative to controls we used the contrasts matrix: (Right brain hemisphere in PD – Left brain hemisphere in PD) – (Right brain hemisphere in controls – Left brain hemisphere in controls). To determine whether hemispheric asymmetry is greater in PD patients or controls, the difference between the absolute fold change of hemispheric asymmetry in PD patients and controls was determined.

To identify CpG/CpH sites exhibiting age-dependent DNA methylation differences in PD patients and controls, we used the contrasts matrix: [(Right brain hemisphere in PD at age max – Left brain hemisphere in PD at age max) – (Right brain hemisphere in PD at age min – Left brain hemisphere in PD at age min)] – [(Right brain hemisphere in controls at age max – Left brain hemisphere in controls at age max) – (Right brain hemisphere in control at age min – Left brain hemisphere in control at age min)]. Age max and min refer to the highest and lowest age value in the cohort, respectively. To determine whether aging changes in hemispheric asymmetry are greater in PD or controls, the absolute fold change of aging differences in hemispheric asymmetry in PD was compared with that of controls.

#### Related to side of symptom dominance (model 2)

DNA methylation changes were examined in PD hemispheres matched or unmatched to side of PD symptom predominance in comparison to control hemispheres. The linear model matrix for CpG/CpH methylation as a dependent variable is:

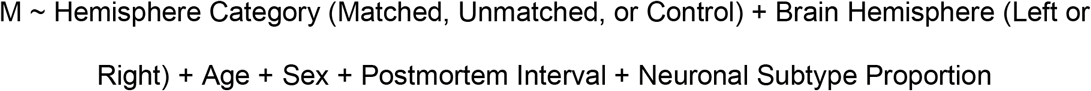

We used the contrasts.fit function to identify cytosine sites exhibiting DNA methylation changes in the matched or unmatched hemisphere of PD patients relative to both hemispheres of controls using the contrasts matrix: Matched PD hemisphere – Control hemisphere, and separately, Unmatched PD hemisphere – Control hemisphere. We identified cytosine sites that were significantly altered in the matched PD hemisphere relative to controls and that exhibited greater DNA methylation fold changes than the unmatched hemisphere.

#### Aging of the symptom-dominant and non-dominant PD hemisphere (model 3)

Aging changes in DNA methylation in the symptom-dominant and non-dominant hemisphere of PD patients were determined. The linear model matrix for CpG/CpH methylation as a dependent variable is:

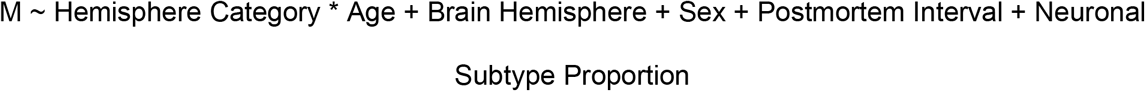

We determined aging changes in hemispheres matched or unmatched to side of symptom dominance using the contrasts matrix: (Matched PD hemisphere at age max – Matched PD hemisphere at age min) – (Unmatched PD hemisphere at age max – Unmatched PD hemisphere at age min).

#### In response to PD disease course (model 4)

Analysis of changes in hemispheric asymmetry of DNA methylation in response to PD disease course was performed using the data from PD patients in the replication cohort, consisting of both hemispheres from the same PD patients with a known side of symptom predominance.

The linear model matrix for CpG/CpH methylation as a dependent variable is:

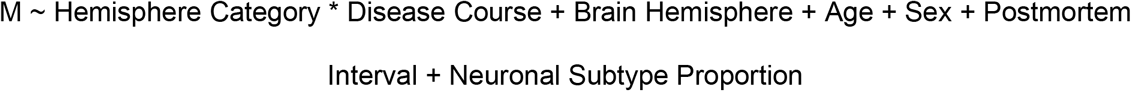

We identified cytosine sites exhibiting changes in hemisphere asymmetry with disease course using the contrasts matrix: (Matched PD hemisphere at duration max – Matched PD hemisphere at duration min) – (Unmatched PD hemisphere at duration max – Unmatched PD hemisphere at duration min). Duration max and min refer to the longest and shortest duration value in the cohort, respectively.

### Gene annotation and enrichment analysis

Because enhancer elements dynamically regulate gene expression through three-dimensional physical interactions, we analyzed chromatin interaction data to reveal the gene targets of enhancers relevant to hemisphere asymmetry. For this analysis we used Hi-C data from the human prefrontal cortex^46^ (Illumina HiSeq 2000 paired-end raw sequence reads; n = 1 sample; 746 million reads; GEO accession ID: GSM2322542). The Hi-C analysis pipeline involved Trim Galore (v0.4.3) for adapter trimming, HiCUP^86^ (v0.5.9) for mapping and performing quality control, and HOMER^87^ (v4.9.1) for identifying significant interactions (default: *p* < 0.001 and zscore > 1.0), with a 40-kb resolution. Hi-C gene annotation involved identifying interactions with gene promoters. This analysis identified a total of 58,758 interactions, genome-wide, of which 23,971 interactions targeted a transcription start site (TSS ± 2 kb). To further identify proximal interactions with gene targets, we used the GREAT^47^ (v4.0.4) software (http://great.stanford.edu/public/html/). Gene annotation was performed for the gene targets of the significant cytosine sites in our analysis and for the background, consisting of gene targets for all cytosines included in our analysis. The background in the discovery dataset was 10,084 genes and in the replication dataset was 10,605 genes.

### RNA-sequencing

We used RNA-seq to profile the mRNA transcriptome in the prefrontal cortex of PD hemispheres with known side of symptom predominance (n = 37 individuals: 13 matched PD hemisphere, 11 unmatched PD hemisphere, and 13 controls). Fresh-frozen human prefrontal cortex tissue (25-50 mg) was lysed and homogenized in 1 mL of TRIzol (Invitrogen) in Precellys Lysing Kit CKMix tubes using a MiniLys homogenizer (Bertin Instruments, 2 intervals of 10 s on the highest setting with 15 s in between). Total RNA was isolated using the standard TRIzol protocol and re-suspended in 85 μL of ultrapure distilled water (Invitrogen). DNase treatment was performed with the RNase-Free DNase kit (Qiagen) using 5 μL DNase I and 10 μL RDD buffer, followed by a column cleanup using the RNeasy Mini kit (Qiagen) with 2 additional washes (75% ethanol) before elution. RNA quantity was assessed by Nanodrop 8000 (Thermo Scientific) and quality was assessed with an Agilent RNA 6000 Nano Kit on a 2100 Bioanalyzer (Agilent Technologies, Inc.). Libraries were prepared by the Van Andel Genomics Core from 500 ng of total RNA using the KAPA RNA HyperPrep Kit with RiboseErase (v1.16) (Kapa Biosystems). RNA was sheared to an average of 300 bp. Prior to PCR amplification, cDNA fragments were ligated to NEXTflex Adapters (Bioo Scientific). The quality and quantity of the finished libraries were assessed using a combination of Agilent DNA High Sensitivity chip (Agilent Technologies Inc.) and QuantiFluor® dsDNA System (Promega Corp.). Individually indexed libraries were pooled, and 75-bp single-end sequencing was performed on an Illumina NextSeq 500 sequencer, with all libraries run across 4 flow cells to return a minimum read depth of 40 million read pairs per library. Base calling was done by Illumina NextSeq Control Software (NCS; v2.0), and the output of NCS was demultiplexed and converted to FastQ format with Illumina Bcl2fastq (v1.9.0).

Trim Galore (v0.11.5) was used to trim the 75-bp single-end reads prior to genome alignment. STAR^88^ (v2.3.5a) index was generated using Ensemble GRCh37.p13 primary assembly genome and the Gencode v19 primary assembly annotation. Read alignment and gene counts were performed using STAR^88^. The gene count matrix was imported into R (v3.5.1) and low expressed genes (counts per million < 1 in all samples) were removed prior to trimmed mean of M-values normalization in edgeR^89^ (v3.16.5). There was 1 sample excluded from further analyses due to poor inter-sample correlations (> 10% difference). At the end of these pre-processing steps, 36 samples with 14,121 genes were retained for downstream analysis. The limma (v3.30.13) statistical package^84^ was used to transform the count matrix to log2-counts per million. A generalized linear model was then used to determine gene transcript level changes in PD hemispheres that are relevant to symptom lateralization (hemisphere category: matched PD, unmatched PD, or control), adjusting for brain hemisphere, age, sex, RIN, and neuron proportion. In addition, we controlled for other sources of variation using RUVSeq^49^ (v1.18.0). We used contrasts to identify differentially expressed genes between matched PD hemisphere *vs.* control as well as unmatched PD hemisphere *vs.* control. *P*-values were adjusted for multiple testing correction using the Benjamini-Hochberg method.

Our RNA-seq analysis is corrected for the proportion of neuronal cells in each sample. Cell-type deconvolution was performed using CIBERSORT^83^ (http://cibersort.stanford.edu), which performs a linear support vector machine learning algorithm on normalized cell type-specific count data. In this approach, we used a gene signature matrix (involving 903 cell-specific marker genes) derived from single-cell RNA-seq measures in adult human brain cells (signature matrix^90^; source^91^). CIBERSORT was run with 100 permutations, and values were used for the neuronal cell composition adjustment in the generalized linear model for transcriptomic analysis.

### DNA methylation status correlated with target gene mRNA levels

We determined whether enhancers and promoters exhibiting DNA methylation changes related to PD symptom lateralization had corresponding changes in target gene transcript levels (n = 36 individuals). For this analysis, we used DNA methylation and RNA-seq data for the same individuals and examined genes exhibiting significant and non-significant DNA methylation changes at their enhancer/promoter in the PD hemisphere matched to symptom dominance. The most significant differentially methylated sites for each target gene were used. For each gene, the diagnosis effect in the RNA-seq data was determined as the residual gene expression in a linear model adjusting for brain hemisphere, age, sex, RIN, neuron subtype proportion, and other sources of variation. For each gene, the diagnosis effect in the DNA methylation was determined as the residual in the linear model adjusting for brain hemisphere, age, sex, postmortem interval, and neuronal subtype proportion. The diagnosis effect in the RNA-seq data was then correlated to that of the DNA methylation data, examining genes with significant (n = 1,208) and non-significant (n = 4,290) DNA methylation changes, separately. Significant association between changes in DNA methylation at gene regulatory elements and changes in corresponding transcript levels in PD (n = 1,208 genes) as compared to non-significant genes (n = 4,290 genes) was determined by the interaction term in the linear regression.

### Pathway enrichment analysis

Pathway enrichment analysis for the genes involved in DNA methylation asymmetry in the healthy brain was done with MetaCore (https://clarivate.com/products/metacore/) and was relative to background genes. Pathway analysis integrating epigenetic and transcriptomic data was performed to identify gene pathways involved in hemispheric asymmetry and symptom lateralization in PD. Pathway analysis was done for DNA methylation data using g:Profiler^92^ and for transcriptomic data was done using GSEA pre-ranked^93^ (v3.0) with Human_GOBP_AllPathways_no_GO_iea_November_01_2017_symbol.gmt from [http://baderlab.org/GeneSets]. Analysis of DNA methylation data identifying pathways involved in hemispheric asymmetry changes with PD disease course was done by g:Profiler^92^. Pathway networks were determined by EnrichmentMap^94^ and annotated by AutoAnnotate^95^ in Cytoscape (v3.7.1). The protein-protein interaction network was performed using STRING^96^ (v11.0).

### Gene ranking

We identified genes having DNA methylation changes most strongly associated with PD symptom lateralization. For this analysis, we used the epigenetic and transcriptomic data from our discovery cohort along with the epigenetic data from our replication cohort. We identified in each dataset the genes preferentially altered in the PD symptom-dominant hemisphere. For DNA methylation data, genes were ranked by p-value of the most significant site. For RNA-seq data, genes in PD were ranked by p-value. We then determined the genes consistently exhibiting greater DNA methylation in symptom-dominant hemisphere of PD patients across DNA methylation and RNA-seq datasets using the aggregateRanks function from the RobustRankAggreg package^97^ (v1.1). We also identified the genes with the most robust aging changes in hemispheric asymmetry in PD patients. As above, genes with significant aging changes in DNA methylation across hemispheres and with differential aging in PD were ranked based on consistency across discovery and replication cohorts, with the most robustly altered genes across datasets determined using the RobustRankAggreg package^97^.

### Mass spectrometry and proteomics analysis

Quantitative proteomic analysis of the prefrontal cortex of PD patients and controls (n = 3 PD and 3 controls) and of the PD prefrontal cortex from hemispheres matched and unmatched to the side of symptom predominance (n = 5 PD-matched and 5 PD-unmatched) was performed. Mass spectrometry analysis of prefrontal cortex samples (∼30 mg tissue) was performed by the Integrated Mass Spectrometry Unit at Michigan State University. Briefly, protein lysate (20 mg) was denatured using 25 mM ammonium bicarbonate/80% acetonitrile and incubated at 37°C for 3 h. The samples were dried and reconstituted in 25 mM ammonium bicarbonate/50% acetonitrile/trypsin solution and incubated overnight at 37°C. The resulting peptides were dried and reconstituted in 25 mM ammonium bicarbonate/4% acetonitrile. Samples were loaded on to a C18 column (2 mm particles, 25 cm x 75 mm ID) and eluted using a 2 h acetonitrile gradient into a Q-Exactive HF-X mass spectrometer. Each sample was run in triplicate to account for technical variance. The mass spectra from each technical replicate were searched against the Uniprot human database using LFQ method in Proteome Discoverer (Version 2.2.0.388, 2017). The technical replicates from each biological sample were pooled and group comparisons (controls vs. PD, PD-matched vs PD-unmatched) were performed using a non-nested test. Only proteins with abundances recorded in at least two samples per group were considered. Proteins with log fold change between groups exceeding ± 0.2 were considered as altered. To identify protein changes in PD relevant to the lateralization of clinical symptoms, we first identified proteins differing between PD and controls, and then merged this protein list with those differing between matched and unmatched PD hemispheres. The resulting list of 345 genes corresponded to disease-relevant proteins involved in hemispheric asymmetry, and their interactions were visualized using STRING-db version 11^96^. Pathway analysis of proteins involved in PD hemispheric asymmetry was done by g:Profiler^92^ with networks determined by EnrichmentMap^94^ and clustered by AutoAnnotate^95^ in Cytoscape v3.7.1.

### Genetic variation associated with DNA methylation

We determined the influence of cis-acting genetic variation on DNA methylation sites relevant to hemispheric asymmetry in PD. BS-SNPer^98^ was used to identify SNPs proximal to the 815,367 cytosine sites profiled in the DNA methylation analysis of the cohort examining both hemispheres of the same PD and control individuals (n = 31 controls and 26 PD patients). SNPs were determined in each sample along with allele frequency by the BS-SNPer software^98^ from bismark generated alignment .bam files. CpG and CpH sites were removed from the list of identified SNPs. SNPs with missing values in more than 50% samples were excluded. The resulting SNPs were assigned to CpG/CpH sites if located within ± 500 kb of the cytosine site. An meQTL analysis was performed examining the effect of genotype on DNA methylation, adjusting for diagnosis, brain hemisphere, sex, age, postmortem interval, and neuronal subtype proportion. A Benjamini-Hochberg false discovery rate correction was performed, with FDR *q* < 0.05 deemed significant. A χ^2^ test was used to compare the number of SNP-methylation associations for cytosines exhibiting hemispheric asymmetry in PD relative to all tested cytosines (background).

